# Cockroach social network position does not influence exploration tendency

**DOI:** 10.1101/2024.04.05.588244

**Authors:** Vasilis Louca, Smilla Savorelli, Nea Vuorio, David N. Fisher

## Abstract

In group-living organisms, we might expect relationships between social position and exploration, such as if certain patterns of social associations encourage exploration or if certain personality types both link different groups and are more likely to explore their environment. However, directly testing these ideas is challenging as exploration can be hard to track in the field and is a difficult trait to measure in the laboratory. Furthermore, invertebrate exploration is less commonly quantified as their small size makes them hard to track. Here we quantified social networks in multiple groups of the gregarious cockroach *Blaptica dubia* in the laboratory and related three measures of social network position (representing overall sociability, connectedness in local cliques, and connectedness to the wider network) to measures of exploration either assayed alone or as part of a group. We found none of the social network measures related to either of the measures of exploration, which were themselves not correlated. We also found that sociability and connectedness to the wider network were positively correlated, suggesting a single axis of variation in centrality within a group. We found no effects of mass or sex, but there were differences between the two blocks our experiments were performed in, suggesting some effect of observer variability. Overall, our results suggest that exploration is not linked to social behaviour in this species, that cockroaches differ in the social behaviour along a syndrome integrating sociability and the tendency to move between groups, and that study replication should be encouraged so we can be confident identified trends are robust.

## Introduction

Organisms explore their environment in order to locate resources such as food, mates, and shelter (Réale et al., 2007; Verbeek et al., 1994). Exploration can influence growth (Adriaenssens & Johnsson, 2011), reproduction (Schuett et al., 2012), and survival (Dingemanse et al., 2004), influencing both lifespan and fitness (Smith & Blumstein, 2008), and therefore plays an important role in ecological and evolution processes. Identifying drivers of exploration and associations with other traits is therefore key for understanding and predicting exploration’s importance and influence in the natural world.

As organisms explore to both avoid others due to competition (Stahl et al., 2001) and find others to mate with (Lipton et al., 2004), social interactions are likely to be key for predicting exploratory tendencies (Louca et al., 2009; Verbeek et al., 1996). Animals engage in social interactions when they mate and cooperate, but also when they fight or compete, and so essentially all animals engage in social interactions of some sort (Székely et al., 2010). The great diversity of social interactions in the natural world, both within and across species, makes it difficult to relate individual social behaviour to other traits such as exploration in a manner that facilitates the synthesis of findings across taxa. A potential solution is to use a common quantitative framework to analyse social interactions and then relate to traits such as exploration. An example of such a framework is social network analysis (SNA; Krause et al., 2007; Wey et al., 2008). In SNA individuals are represented as “nodes” or “vertices” and are connected with other individuals with “ties”, “edges”, or “links” (see: Croft et al., 2008; and: Krause et al., 2014 for introductions). This approach is represented mathematically as a matrix, where positive valued cells are present between individuals when they interact and 0 values are used when they do not. We can then use the extensive suite of measures in SNA (Sosa et al., 2021) to quantify different aspects of social behaviour, including those depending on both direct and indirect interactions (Brent, 2015), and relate to other traits and compare across species (Farine & Whitehead, 2015; e.g.: Sueur et al., 2011).

Testing the link between social behaviours and other behaviour traits such as exploration is especially important in group-living species, as in such taxa social interactions are near-constant and so could strongly impact other traits. It is conceivable that, in group living animals, social interactions are related to exploration in various ways. Negative interactions may stimulate an individual to leave their current group or area (e.g. in the western bluebird *Sialia mexicana*; Aguillon & Duckworth, 2015; Christian, 1970). In contrast, positive associations may encourage an individual to stay (the “social cohesion hypothesis”; Bekoff, 1977; e.g. in yellow-bellied marmots *Marmota flaviventris*; Blumstein et al., 2009).

Meanwhile, an organism’s own social tendencies might be related to its exploration tendency, for instance unsociable individuals might be more willing to explore for new groups to join rather than remain in groups of high densities, while sociable individuals might be more willing to search for new groups to join when at low densities (as seen in the common lizard *Lacerta vivipara*; Cote & Clobert, 2007). In the males of the spotted hyena (*Crocuta crocuta*), there is evidence that during ontogeny, network connections tend to weaken, which eventually leads to individuals dispersing from their natal groups (Turner et al., 2018). These relationships may differ between the sexes, for example if females compete for resources they be more exploratory at high density, while if males compete for access to females they may be more exploratory at low densities (as seen in feral horses *Equus ferus caballus*; Marjamäki et al., 2013). There may also be confounding factors for these relationships, for instance higher body condition may effect position in a social group (for example in female reindeer *Rangifer tarandus*; Holand et al., 2004) but also tendency to explore (such as in zebra finches *Taeniopygia guttata*; Crino et al., 2017), creating the illusion of a relationship between social phenotype and exploration when in fact none exists. Explicitly testing the relationship between exploration and social behaviour, and how it depends on context, is therefore essential. This is especially true in invertebrates, as their exploration is relatively understudied (Kralj-Fišer & Schuett, 2014).

When relating exploration to social behaviours, it could be important to quantify exploration in both solitary and group contexts. As noted above, exploration might directly depend on social interactions. Additionally, the motivation to stay within a social group, or coercion to either stay or leave, might mask variation in innate tendency to explore, which would only be revealed when testing individuals alone. Finally, exploration can be inconsistent across contexts (Arvidsson et al., 2017), and tests behavioural ecologists perceive to be assessing the same trait may in fact not be (Carter et al., 2013). It is therefore useful to quantify exploration in multiple situations and determine if the behaviour is correlated across context.

Here we develop methods to quantify exploration tendency both when alone and in group contexts. We relate each measure of exploration to three measures of social behaviour based on social networks in a laboratory population of the cockroach *Blaptica dubia*. We also compare the sexes and control for body mass. This allows us to measure how different aspects of social behaviour relate to exploration and how these traits are affected by sex while controlling for key confounding variables.

We predict that individuals with a higher exploratory tendency will have a higher strength (a measure of gregariousness), as they will be moving regularly between subgroups. For the same reason we expect high exploration to be associated with lower clustering coefficient (a measure of how frequently an individual’s associates themselves interact), as individuals with connections in different subgroups will not have associations that are themselves connected. Finally, we expect higher exploration to be associated with higher closeness (a measure of connectedness across the entire network), as individuals moving between subgroups will be well-connected to the wider network. We also predict that males, which unlike females in this species have fully developed wings (but with limited ability to fly; Kesel et al., 2009) and are likely the dispersing sex, will show higher exploratory tendencies. Body condition could affect exploration in either direction; better condition individuals may have the stored resources to take on the risk of exploring, but alternatively poor condition individuals may explore to find more resources (Luttbeg & Sih, 2010). As such, we do not make a directional prediction.

## Methods

### Study population

A population of *B. dubia* was founded in the lab in Aberdeen in March 2021 by purchasing a mixed colony online. They were then maintained at 28°C, 50% humidity, and a 12:12 light:dark cycle. Stock was maintained in 48L plastic boxes (610 × 402 × 315 mm), with cardboard egg tray as shelter, carrot for hydration and Sainsbury’s dog food for nutrition (approx. nutritional composition = 1527 kJ energy, 24 g protein, 12 g fat per 100 g). Boxes of immature nymphs were of mixed ages and sexes, with individuals that eclosed into adults moved to adult-only boxes within three-four days of eclosion. Some adult boxes were used for breeding and so contained both sexes, while others were for experimental animals and were single-sex. The colony showed no infection in this time and low mortality in all life stages, and so presumed to be healthy (for further details on the stock population see: Fisher, 2023). Data collection took place in two blocks with one researcher conducting the majority of the work in each, the first from 26/5/22 to 14/7/22 (led by SS), the second from 7/10/22 to 14/11/22 (led by NV). The methods were the same in each block unless we note otherwise.

### Data collection

We collected data in each block in three stages, first measuring social associations to construct social networks, then assaying the cockroaches’ exploration in a group setting, before measuring exploration alone. To create groups to quantify social networks, we selected unmated adult males and females from single-sex stock boxes and placed them into single-sex groups of ten (selection of individuals was not random, as that would have required labelling each individual in the stock population, but we removed shelters from boxes before selecting individuals to avoid only selecting “bolder” individuals). We created 12 groups in the first block (six of each sex) and ten in the second block (five of each sex). The high level of replication of networks (22 in total) is a particular strength of our study; while most studies on social networks in animals only have a single or few networks, we have multiple independent networks. This replication prevents the behaviour of a few individuals influencing the social phenotypes of every individual in the study (as social network measures often depend on connections in across the whole network). We weighed each individual and, so that we could recognise individuals, for the first block marked each with a unique combination of two colours (red, green, blue, white and gold) on their wing cases using paint pens (Edding 780 Extra-fine paint markers). As we filmed the second block in darkness, here we instead marked individuals with a unique combination of dot locations (dots present/absent on head, left wing case and right wing case, with the head occasionally having 2 dots to reach a unique code for all 10 individuals) using only white paint pens. We placed each group of ten into a plastic box (340 × 200 × 125 mm) with four cardboard egg-tray shelters (approx. 100 x 120 mm), one in each corner taped along the long wall so they are not touching, creating four separate shelters (see: Fisher, 2023 Fig. 1b). We placed dog food (enough to ensure cockroaches have excess food, around 3-5g) and carrot (for hydration, around 10g) in each box between the shelters and replaced it each time we recorded associations. To record associations, we observed which individuals were using the same shelter on six dates between two to four days apart (from 26/5/22 to 13/6/22 in the first block and from 7/10/22 to 19/10/22 in the second). Observations were only roughly evenly spaced due to avoiding data collection at weekends and other occasional scheduling constraints, but observations never took place on consecutive days. After recording the associations of all individuals in a box they were removed from the shelters and placed in the centre of the box, so any associations would be re-formed each observation. In this manner we gain six snapshots of the social associations in each group of cockroaches, much like observations of flocks of birds or herds of ungulates. We then aggregated these data to build a single social network for each group by assuming those in the same shelter are associated in some way (the “gambit of the group”; Whitehead & Dufault, 1999). While a single observation of two individuals sharing a shelter does not imply they are close social associates, consistently observing them together would allow us to infer some degree of social preference. Using the R package *asnipe* (Farine, 2013), we converted the data on co-occurrence in groups into an undirected weighted social network, where edge weights equalled the “simple ratio index”; the number of times the two individuals were using the same shelter divided by the total number of times they were seen (Cairns & Schwager, 1987). This value is 1 if two individuals are always seen together and near zero if they are very rarely seen together (it is exactly zero if they are never seen together). We did this separately for each group, giving 22 independent social networks, 11 of each sex.

**Figure 1.**
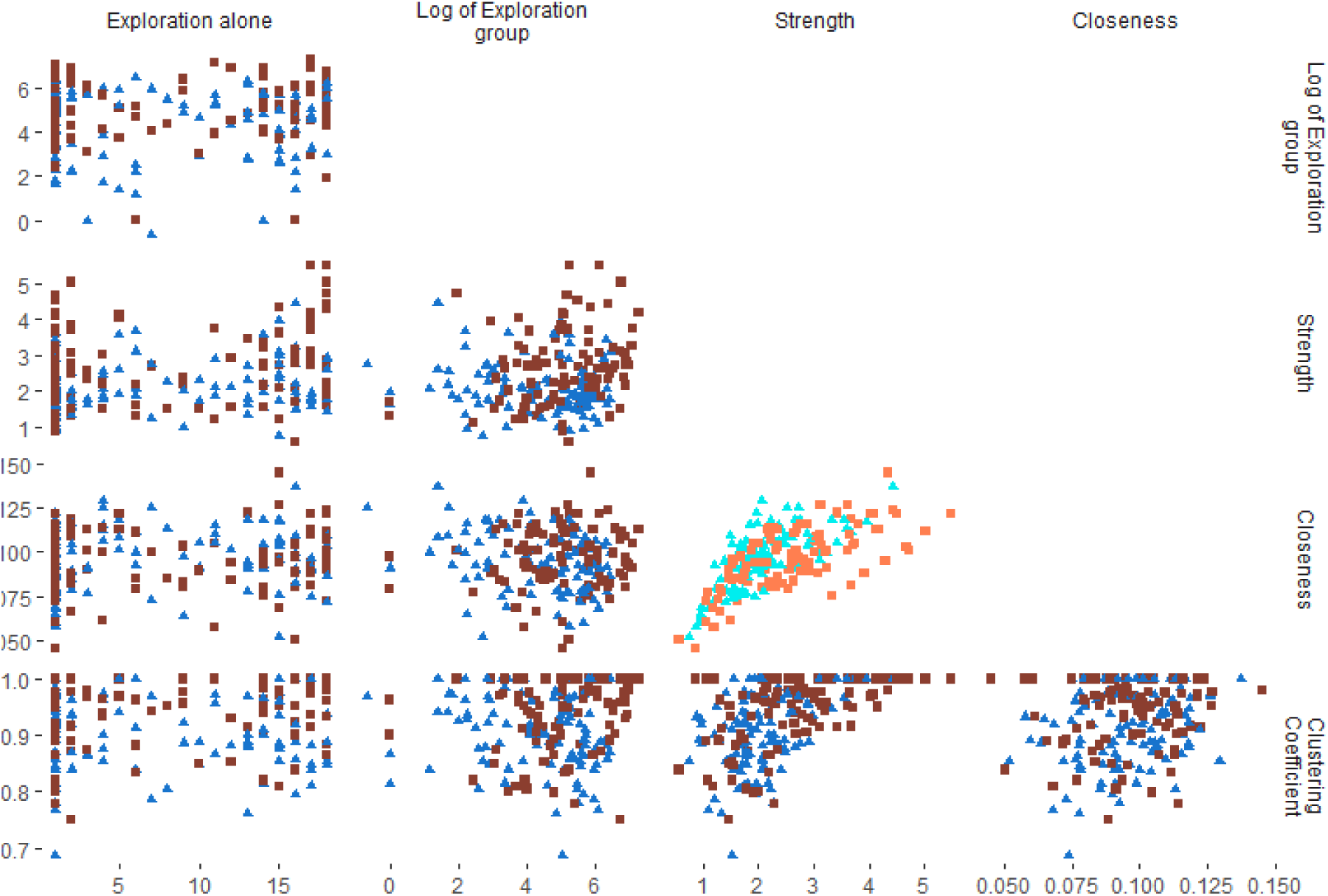
A pairs plot showing the relationship between each pair of traits. Females are orange squares, males are blue triangles, with the statistically significant correlation (closeness – strength) highlighted in lighter colours.

For each individual we calculated three social network measures, each representing a different aspect of social behaviour. These measures all are weighted, and hence account for the frequency of the associations. First was “strength”, the sum of all an individual’s weighted interactions, representing overall gregariousness (function **rowSums** in the package *Matrix*). Second was “clustering coefficient”, the sum of the weighted edges involved in *closed* triads centred on the focal node divided by the sum of all weighted edges in triads centred on the focal node (Barrat et al., 2004; Opsahl & Panzarasa, 2009), which represents whether individuals invest in a closed circle of associates by regularly interacting with others who they share a mutual association with or not (function **clustering_local_w** in *tnet*; Opsahl, 2009). Finally, there was “closeness”, which is the inverse of the mean of the weighted pathlengths between an individual and all others in the network (Freeman, 1978), representing how well connected to all parts of the group the individual is (function **closeness_w** in *tnet* Opsahl, 2009; Opsahl et al., 2010 using the standard [non-normalised] measure, since all our networks are 10 individuals).

After the sixth observation of associations, individuals in block 1 were immediately moved to the next stage of data collection. However, individuals in block 2 were observed a further four times (giving 10 in total). To better compare the blocks, we do not use the extra four observations in block 2 here (social network measures from networks based on six observations are either strongly or moderately positively correlated with measures from networks based on ten observations; Supplementary Materials Fig. S1). In the next stage we tested the individuals for exploratory tendency in group setting (hereafter “exploration-group”). We removed the carrot from the groups for 24 hours in block 1 and 48 hours in block 2 to increase their need for hydration and so willingness to explore; the change between groups was made to encourage more individuals to explore their environment (although this was unsuccessful as there was no difference in mean exploration-group latency between the blocks, see Results). We then transferred all group members into a single large plastic box (610 × 402 × 315 mm) with the four shelters used in the social association stage placed at one end of the box, evenly spaced. At the other end of the box we placed 10g of carrot and three pellets of dry dog food (Sainsbury’s Complete Nutrition Adult Small Dog Dry Dog Food, approx. nutritional composition = 1527 kJ energy, 24g protein, 12g fat per 100g) to act as a goal. We placed each box in a room maintained at 22-26°C (although with no humidity control), with a pet heating pad placed beneath each box to raise the temperature closer to that of the stock room to encourage movement (22-24^0^ C). Above each box, at approximately 50cm, we placed an IP camera (ABUS IP video surveillance 8MPx mini tube camera) pointing directly downwards to record the box from above. We then left the room for 24 hours, with the camera recording continuously. For block 1, the lights were left on, while for block 2 the lights were turned off as, similarly to the above-mentioned difference in carrot deprivation between groups, this was to encourage more individuals to leave the shelter, although as noted above this was unsuccessful.

We were able to record up to four groups simultaneously, and so we filmed them in batches of two to four (which means groups had different periods between the end of the recording of the social associations and the start of the exploration-group trial). In block 1 we initially made recordings on 14-23/6/22, but due to computer error these were not useable, and so we repeated the trials on 1-6/7/22 (therefore starting at least 22 days after the final observation of social associations) and only used the data from the second run. In block 2 the recordings were made from 7-9/11/22 (therefore starting at least 18 days after the final observation of social associations). From the video recordings we extracted the number of minutes between the start of the trial and when each individual first crossed the half-way point of the box (i.e., it left the shelter and travelled a distance of approx. 200 mm). This measure of exploration is therefore low when individuals are quick to explore, and high when they are slow/unwilling to explore (with a theoretical maximum of 1440). We gave individuals who never crossed the half-way point values of NA to avoid a data distribution with a spike at high values (n = 11). We also repeated the analysis with those that did not explore given a score of 1440; the clear results in the original analysis are maintained in the second analysis with the additional presence of a mass-exploration group relationship in males (see Supplementary Materials). After 24 hours we returned each group to the box it had recorded associations in with fresh carrot and cardboard shelters and returned them to the stock room.

For the third stage we measured a proxy for exploratory tendency while alone (hereafter “exploration-alone”) by placing each individual in a small plastic box (200 × 100 × 70 mm) with a grid of 18 rectangles (each 32 by 27 mm) drawn onto paper underneath the box’s (clear) floor. In the same room as we recorded exploration-group, again using pet heating pads to raise the temperature, we placed boxes with cockroaches in sets of four beneath the same IP cameras, switched off the lights (as *B. dubia* is more active in the dark; Bouchebti et al., 2022), and filmed continuously for 30 minutes. These cameras automatically use infrared LEDs to film in the dark, which we presume the cockroaches cannot see. We ignored the first 10 minutes of the recording as we did not want to record the individual’s immediate response to being placed in a box, and instead counted the number of unique squares each individual visited in 20 minutes as a measure of exploratory tendency (counting unique areas visited as opposed to total amount of movement separates this from a measure of general activity ; Réale et al., 2007). In block 1 we made recordings on 12-14/7/22 (therefore starting at least 7 days after assaying exploration-group), while in block 2 we made recordings on 10-14/11/22 (with trials staggered so that individuals were tested 3 or 4 days after being assayed for exploration-group). After the exploration-alone trials we returned individuals to the stock population. Individuals that died part way through the experiment had the values for the behavioural trials they completed retained in the dataset and NA values entered for any missing (n = 7).

### Data analysis

Our final dataset comprised of 215 individuals measured for each of the three social network traits, 199 of which recorded a non-NA value for exploration-group, and 208 which recorded a measure of exploration-alone. To determine how the different behaviours were associated, and if any were influenced by mass or sex, we entered the five behaviours into a multivariate mixed-effects model using function **MCMCglmm** in the R package *MCMCglmm* (Hadfield, 2010). We used each social network trait, exploration-alone, and the log of exploration-group as response variables and estimated the covariances among each of them simultaneously (10 covariances in total). We included block number and sex as two-level categorical variables, and cockroach mass as a continuous predictor, mean centred and scaled to a variance of 1 (Schielzeth, 2010). We also included the interaction between (scaled) mass and sex to test whether the effect of mass differed between the sexes. We estimated an intercept and the relationship with each fixed effect for each response variable, giving 25 coefficients in total. We included the random effect of group identity, to estimate any among-group variation that may have arisen due to emergent group dynamics and to account for differences in what the groups experiences e.g., the slightly different periods of time between behavioural tests. We did not estimate the among-group covariance in behavioural traits, leaving the covariance as the phenotypic covariance, once the effect of the fixed effects and the variation in means among groups was accounted for.

We ran the model for 550,000 iterations, discarding the first 50,000 (as parameter estimates change most at the start of the chain), and using 1 in each 100 iterations to generate the posterior distributions. We used a very weakly informative prior with a variance of 1 and a degree of belief of 0.002 for the random effect and residual variance. For exploration-alone we used a Poison error distribution, while for the other four traits we used a Gaussian error distribution as the raw data distributions were not too far from normality and we achieved satisfactory convergence with these error distributions. We calculated the among-trait correlations as the covariance divided by the square root of the product of the variances of the two traits. We did this for each draw in our posterior distribution, allowing us to estimate the mode and 95% credible intervals for each correlation; those with credible intervals that do not overlap zero are considered clearly different from zero. For the fixed effects we used the posterior distributions; those with 95% credible intervals that did not overlap zero have a relationship with the trait in question. The author that ran the analysis (DNF) did not conduct the behavioural trials or extract data from the video, and so was blind to any impressions developed during those stages.

## Results

Closeness was very strongly positively associated with strength (R = 0.939, CrIs = 0.918 to 0.953; we show the raw correlations in Fig. 1 and the correlation posterior distribution modes and the 95% credible intervals estimated from the model in Fig. 2). Clustering coefficient was not associated with closeness (R = -0.001, CrIs = -0.137 to 0.152) or strength (R = 0.002, CrIs = -0.138 to 0.148). Neither exploration-group nor exploration-alone were associated with any of the social network traits (exploration-alone – strength R = 0.046, CrIs = -0.120 to 0.046; exploration-alone – clustering coefficient R = -0.017, CrIs = -0.182 to 0.154; exploration-alone – closeness R = 0.070, CrIs = -0.089 to 0.227; exploration-group – strength R = -0.058, CrIs = -0.219 to 0.079; exploration-group – clustering coefficient R = - 0.067, CrIs = -0.245 to 0.053; exploration-group – closeness R = -0.093, CrIs = -0.228 to 0.066). The two measures of exploration were not associated with each other (R = -0.104, CrIs = -0.242 to 0.089). The residual variances and covariances are given in Table S1.

**Figure 2.**
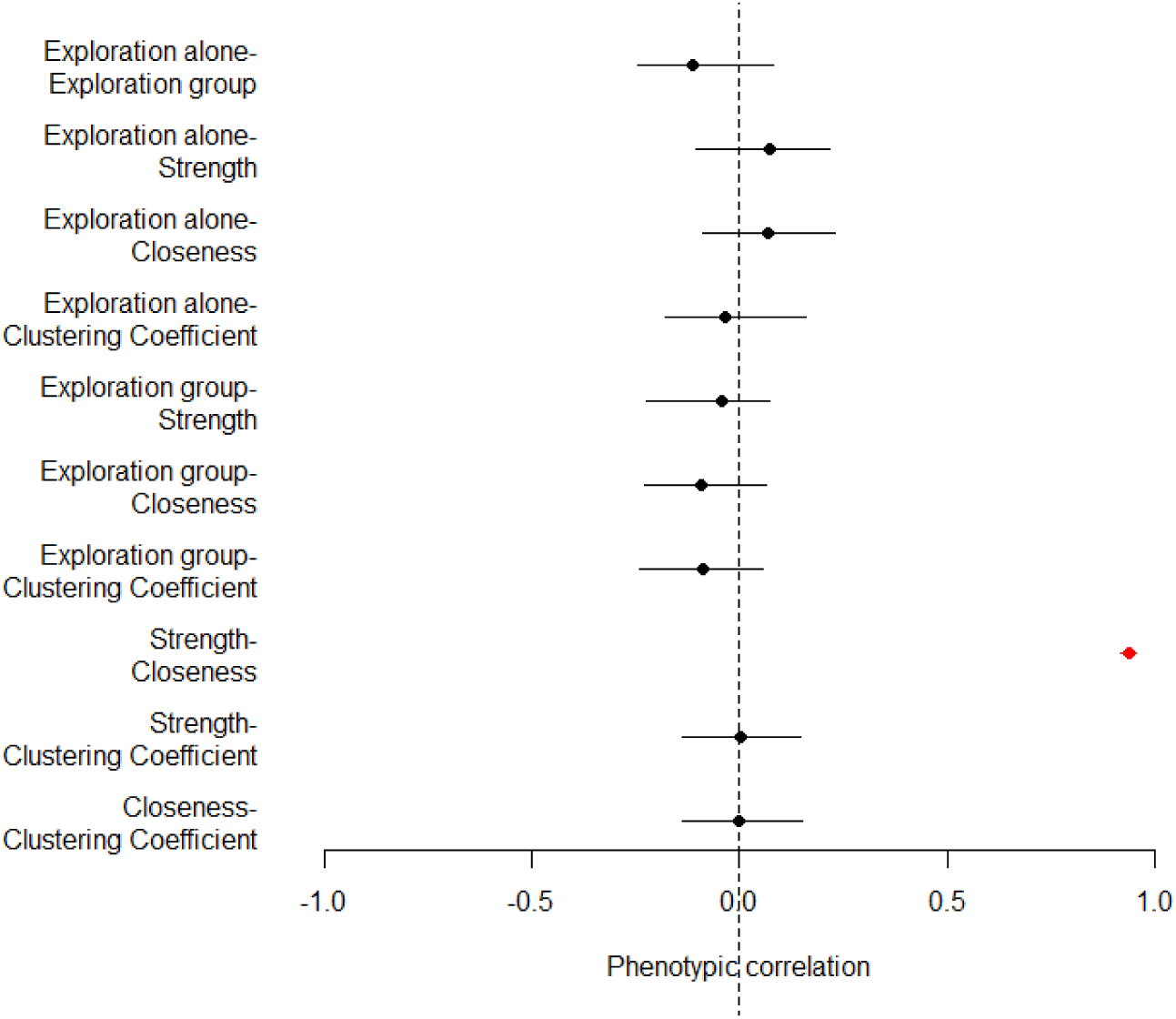
Correlations between traits as estimated from the model., Points show the posterior distribution mode, the horizontal lines show the 95% credible intervals. The correlation that did not overlap with zero has been highlighted in red.

Cockroach mass was not associated with any behavioural trait in either sex. Males had marginally lower strength scores, although this latter effect overlapped with zero (Fig. 1). The sexes did not differ in any other trait. Individuals in block 1 had lower strengths and exploration-alone scores than individuals in block 2 (model results in Table 1).

**Table 1.**
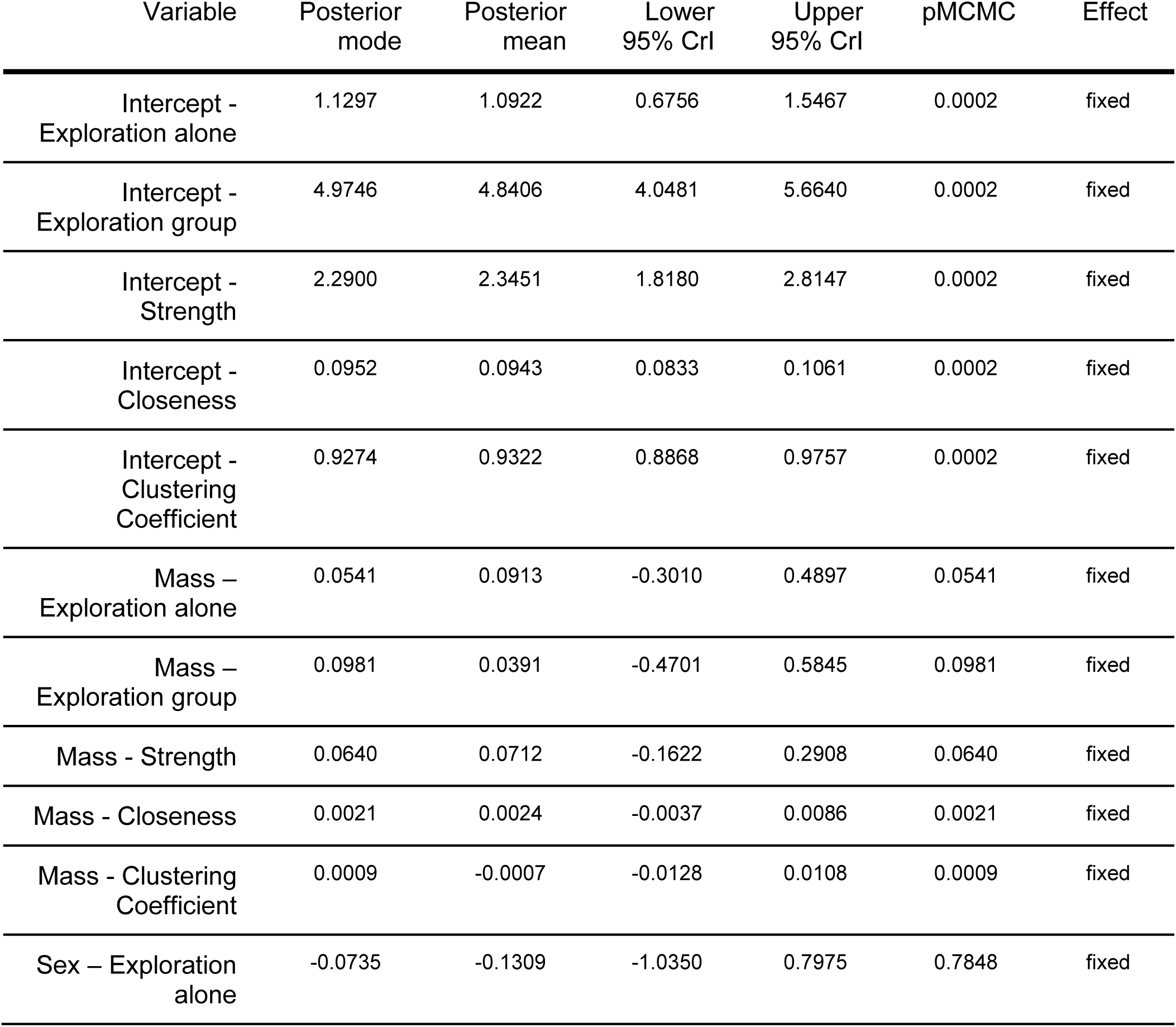
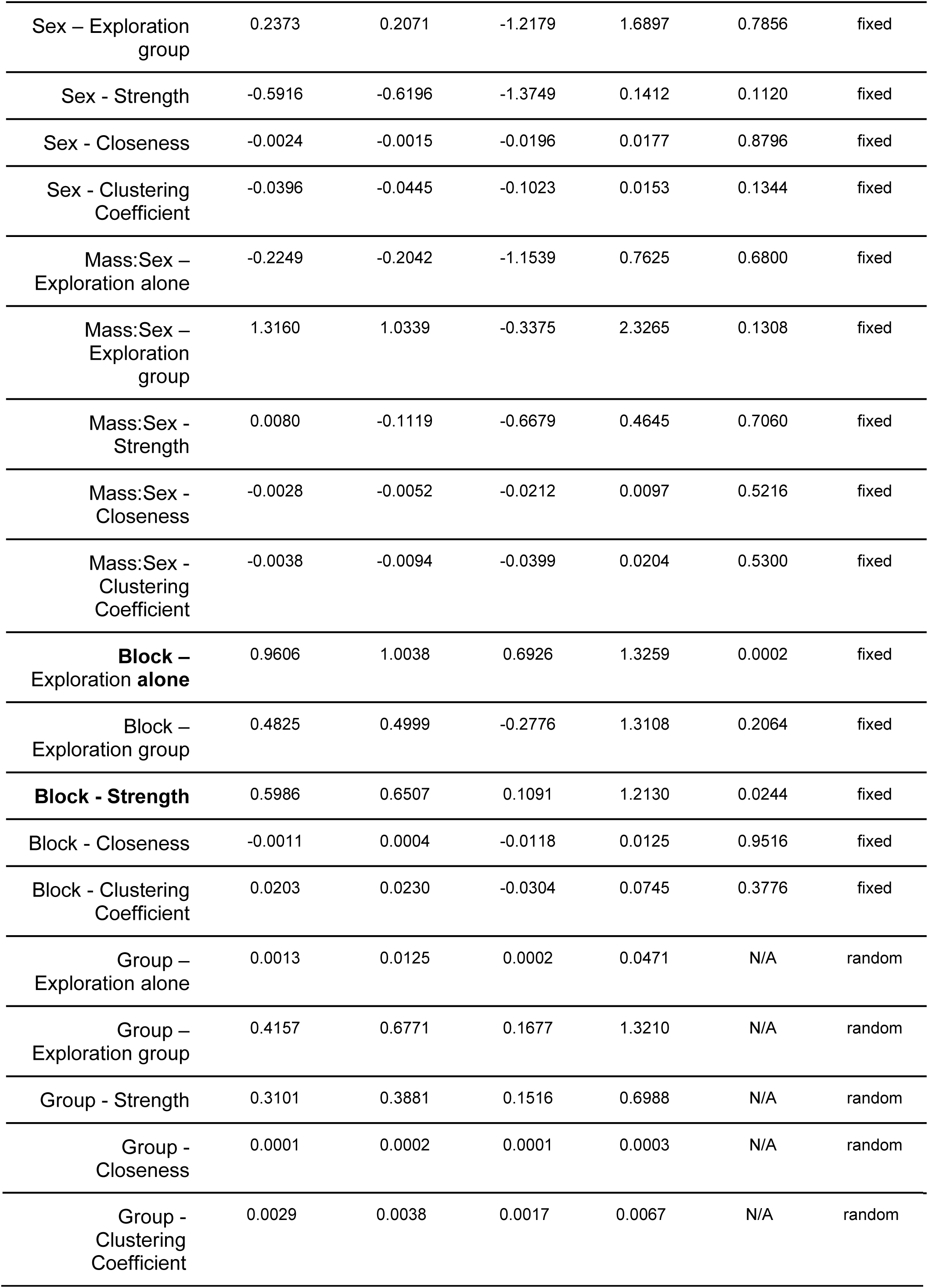
Additional effect estimates from our statistical model. For each of the five traits (exploration alone, exploration group, strength, closeness, and clustering coefficient) we give the intercept, the effect of mass, the effect of sex (females are the default, so this effect represents how males deviate from females), the mass:sex interaction (again, as females are the default this effect indicates how the effect of mass on male traits differs from that for females, which is indicated by the main effect of mass), the effect of block (block 1 is the default, and so this effect indicates how block 2 differs), and the random effect of group. As group is a random effect, we only report the posterior mode, mean, and lower and upper 95% credible intervals (CrI), while as the remaining terms are fixed effects we also report the pMCMC value, which represents two times the probability the effect is either greater or smaller than zero (whichever is smaller), and can be interpreted in a similar way to traditional (frequentist) p-values. Terms where the 95% CrIs do not overlap with zero are highlighted in bold.

## Discussion

Our study investigated aspects of individual social network position within groups of *Blaptica dubia* cockroaches and demonstrated that different measures of exploration are not linked to these positions. Two of the social network measures, strength and closeness coefficient, were very strongly positively correlated with each other, but neither correlated with a third, clustering coefficient. Meanwhile, our two measures of exploration were not correlated. Therefore, while we had predicted social interactions may be associated with exploration (with the direction of causality plausibly in either direction, or both being affected by a third variable) we found no evidence for this prediction. Instead, exploration in *B. dubia* might depend on other aspects of behaviour, such as the motivation to mate and reproduce (which since all our individuals were unmated did not vary in this study), such that the variables we did measure were unimportant.

Cockroach mass was not found to influence exploration (or social network position) in either sex, demonstrating either that animal condition is not a driver for exploration or that mass is not a good proxy for body condition in our system. There is evidence that in German cockroaches (*Blattella germanica*) exploration is linked to food availability and starvation (Ross & Tignor, 1985; Tignor & Ross, 1987), which might indicate that in some cockroach populations lack of food and hunger might initially be a driver for exploration, but in a stable laboratory population with ample access to resources such as ours, there is not sufficient pressure on the lighter/poorer condition individuals to explore. We could test this by experimentally limiting food availability, beyond the 48 hours we used in block 2 of this study. This manipulation would also be instructive for understanding the relationship between mass/condition and social behaviour, as we found no relationship here. Studies on some species, such as sleepy lizards (*Tiliqua rugosa*) and great tits (*Parus major*), have also suggested limited relationship between body condition and social network structure (Godfrey et al., 2013; Snijders et al., 2014), but other studies have suggested relationships between connectedness and body mass (e.g., in prairie voles *Microtus ochrogaster*; Sabol et al., 2020). As the number of studies using social network analysis grows, we may be able to integrate across studies (either informally or formally as a meta-analysis) to determine the typical relationship between mass/condition and centrality measures, and if this relationship varies systematically across certain categories (such as between the sexes) or groups (such as among taxa) of animals.

Perhaps the most surprising result of our study is the lack of a clear relationship between our two measures of exploration. We had designed our assays to capture exploratory tendency in two different contexts, but it is possible that instead we measured two different traits. One possibility is that exploration alone, which was quantified as searching a small arena, functions to locate food or water, while exploration in a group, which was assayed in a larger arena, functions as a form of dispersal to locate mating partners. Along these lines, movement over longer distances in the German cockroach (*Bl. germanica*) is often linked to searching for mates for breeding (Ross & Tignor, 1985), which may assist with avoiding inbreeding (Lihoreau & Rivault, 2010). In contrast, exploring the more immediate area may serve to search for food, rather than dispersing to search for mating partners (Durier & Rivault, 2001). Despite the different functions, we might still have expected such similar behaviours to be positively correlated, such as if individuals with a higher metabolic rate both moved around their immediate environment more and travelled long distances more (Biro & Stamps, 2010). In other species positive correlations are seen, such as in roe deer (*Capreolus capreolus*) and a colonial spider (*Cyrtophora citricola*), where different measures of exploration (exploration beyond the home range and natal dispersal, and exploration of a new environment and “tiptoeing” to release silk into air, respectively) are positively correlated (Debeffe et al., 2013; Yip et al., 2021, p. 202). As Dochtermann (2023) highlights, we still have a limited understanding of when and how different behavioural traits covary. Therefore, until we generate better theoretical predictions, we will have to sift through published examples and try to detect broader patterns.

A not-mutually-exclusive alternative to these measures of exploration being different behaviours is that exploration shows such high within-individual variability that a correlation across the two contexts (as well as across the time between the assays) was too weak to be detected. Some personality traits in cockroaches are consistent (Arican et al., 2020; Planas-Sitja & Deneubourg, 2018; Stanley et al., 2017), suggesting we might expect exploratory behaviours to be consistent too. However, further data on cockroach exploration in controlled conditions (as opposed to in uncontrolled settings such as through apartment buildings; Barcay et al., 1990; Crissman et al., 2010) is lacking for us to suggest why in our study we might not have found this. In great tits (*P. major*) exploration of a novel environment is repeatable and heritable (Dingemanse et al., 2002), and genetically correlated with exploration over a wider area (dispersal distance), yet the phenotypic correlation between these two traits is not different from zero (Korsten et al., 2013), or is only positive for females in some seasons, and never in males (van Overveld et al., 2014). In the case of the great tits, a negative environmental covariance counters the positive genetic one to give the phenotypic null covariance (Korsten et al., 2013); the same might be true in our system, but we would need further work on the genetics underpinning cockroach behaviour to find out.

We found that strength was strongly positively correlated with closeness. The correlation is so strong (R = 0.939) that these represent essentially the same trait: cockroaches that are highly gregarious are well connected to all parts of the network. This result suggests individuals differ in their social behaviour from less gregarious/central in the network to more gregarious/central. Planas-Sitja and Deneubourg (2018) found that collective aggregation formation in the American cockroach (*Periplaneta americana*) is influenced by the social behaviour of individuals in the aggregations, suggesting substantial differences among-individuals. Such differences in social behaviour will likely expose an individual to a range of different selection pressures, such as differential exposure to directly transmitted infections, variation in competition for resources and mates, and different risk of desiccation. Understanding what drives this variation in social behaviour and what consequences it has for the ecology and evolution of cockroach groups are important next steps.

We also found that males and females had similar social network positions. This is in contrast to work in another Blaberid, *Diploptera punctata*, which found that females were more gregarious and formed the core of a proximity network (Stanley et al., 2018). We did not assay male and females together in a social network, whereas Stanley et al. did, which may explain the difference. In mixed-sex groups, females may be more central as they are larger, and so more able to displace others from preferred spots in an aggregation, or as they are preferred associates of both males and other females, causing them to end up in the centre of an aggregation (Stanley et al., 2018). In single-sex groups, such inter-sex interactions are impossible and so social networks will predominantly reflect individual social tendencies. Measuring social associations in mixed-sex groups in *B. dubia* would allow us to more directly compare our results to those of Stanley et al. and determine whether the differences are due to experimental design or a difference in behaviour between these two species.

Our study has also demonstrated that two traits (exploration alone and strength) differed between blocks, which indicates the presence of observer-induced variation. This could come about through the methodological differences between the blocks we have already noted, or due to subtle differences in the way the two researchers leading blocks 1 and 2 (SS and NV respectively) handled the cockroaches. That the mean level of a behaviour that is observed differs between blocks within a single study should be considered in other studies and when making conclusions from single studies at single point in time by a single laboratory, reinforcing the need for more extensive replication in such experiments (Brecht et al., 2021; Farrar et al., 2021; Gaona-Gordillo et al., 2023).

In conclusion, our results suggest care should be taken when measuring exploration in the laboratory, as assays designed to capture exploration may actually measure different traits or not capture the consistent component of exploration. We also found no support for the suggestion that social behaviour and exploration are associated. We found limited importance of mass or sex in our system, while there was a single axis of variation from more central to less central individuals, possibly reflecting individual variation in social behaviour. Finally, our results indicate that repeating experiments, even in the same lab, can give different results, and so add strength to the call that study replication should be an important component of animal behaviour research if we are to detect general pattens.

## Acknowledgements

We thank Keith Lockhart for maintaining the cockroach stock population.

## References

Adriaenssens, B., & Johnsson, J. I. (2011). Shy trout grow faster: Exploring links between personality and fitness-related traits in the wild. Behavioral Ecology, 22(1), 135–143. 10.1093/beheco/arq185

Aguillon, S. M., & Duckworth, R. A. (2015). Kin aggression and resource availability influence phenotype-dependent dispersal in a passerine bird. Behavioral Ecology and Sociobiology, 69(4), 625–633. 10.1007/s00265-015-1873-5

Arican, C., Bulk, J., Deisig, N., & Nawrot, M. P. (2020). Cockroaches Show Individuality in Learning and Memory During Classical and Operant Conditioning. Frontiers in Physiology, 10, 1539. 10.3389/FPHYS.2019.01539/BIBTEX

Arvidsson, L. K., Adriaensen, F., van Dongen, S., De Stobbeleere, N., & Matthysen, E. (2017). Exploration behaviour in a different light: Testing cross-context consistency of a common personality trait. Animal Behaviour, 123, 151–158. 10.1016/j.anbehav.2016.09.005

Barcay, S. J., Schneider, B. M., & Bennett, G. W. (1990). Influence of Inseċticide Treatment on German Cockroach (Dictyoptera: Blattellidae) Movement and Dispersal within Apartments. Journal of Economic Entomology, 83(1), 142–147. 10.1093/jee/83.1.142

Barrat, A., Barthélemy, M., Pastor-Satorras, R., & Vespignani, A. (2004). The architecture of complex weighted networks. Proceedings of the National Academy of Sciences of the United States of America, 101(11), 3747–3752. 10.1073/pnas.0400087101

Bekoff, M. (1977). Mammalian Dispersal and the Ontogeny of Individual Behavioral Phenotypes. The American Naturalist, 111(980), 715–732. 10.1086/283201

Biro, P. A., & Stamps, J. A. (2010). Do consistent individual differences in metabolic rate promote consistent individual differences in behavior? Trends in Ecology & Evolution, 25(11), 653–659. 10.1016/j.tree.2010.08.003

Blumstein, D. T., Wey, T. W., & Tang, K. (2009). A test of the social cohesion hypothesis: Interactive female marmots remain at home. Proceedings of the Royal Society of London B: Biological Sciences, 276(1669), 3007–3012.

Bouchebti, S., Cortés-Fossati, F., Estepa, Á. V., Lozano, M. P., Calovi, D. S., & Arganda, S. (2022). Sex-Specific Effect of the Dietary Protein to Carbohydrate Ratio on Personality in the Dubia Cockroach. Insects 2022, *Vol.* 13, *Page* 133, *13*(2), 133. 10.3390/INSECTS13020133

Brecht, K. F., Legg, E. W., Nawroth, C., Fraser, H., & Ostojic, L. (2021). The Status and Value of Replications in Animal Behavior Science. Animal Behavior and Cognition, 8(2), 97–106. 10.26451/abc.08.02.01.2021

Brent, L. J. N. (2015). Friends of friends: Are indirect connections in social networks important to animal behaviour? Animal Behaviour. 10.1016/j.anbehav.2015.01.020

Cairns, S. J., & Schwager, S. J. (1987). A comparison of association indices. Animal Behaviour, 35(5), 1454–1469. 10.1016/S0003-3472(87)80018-0

Carter, A. J., Feeney, W. E., Marshall, H. H., Cowlishaw, G., & Heinsohn, R. (2013). Animal personality: What are behavioural ecologists measuring? Biological Reviews, 88(2), 465–475. 10.1111/brv.12007

Christian, J. J. (1970). Social Subordination, Population Density, and Mammalian Evolution. Science, 168(3927), 84–90. 10.1126/science.168.3927.84

Cote, J., & Clobert, J. (2007). Social personalities influence natal dispersal in a lizard. Proceedings of the Royal Society B: Biological Sciences, 274(1608), 383–390. 10.1098/rspb.2006.3734

Crino, O. L., Buchanan, K. L., Trompf, L., Mainwaring, M. C., & Griffith, S. C. (2017). Stress reactivity, condition, and foraging behavior in zebra finches: Effects on boldness, exploration, and sociality. General and Comparative Endocrinology, 244, 101–107. 10.1016/j.ygcen.2016.01.014

Crissman, J. R., Booth, W., Santangelo, R. G., Mukha, D. V., Vargo, E. L., & Schal, C. (2010). Population Genetic Structure of the German Cockroach (Blattodea: Blattellidae) in Apartment Buildings. Journal of Medical Entomology, 47(4), 553–564. 10.1603/me09036

Croft, D. P., James, R., & Krause, J. (2008). Exploring Animal Social Networks. Princeton University Press. http://opus.bath.ac.uk/25878/

Debeffe, L., Morellet, N., Cargnelutti, B., Lourtet, B., Coulon, A., Gaillard, J. M., Bon, R., & Hewison, A. J. M. (2013). Exploration as a key component of natal dispersal: Dispersers explore more than philopatric individuals in roe deer. Animal Behaviour, 86(1), 143–151. 10.1016/j.anbehav.2013.05.005

Dingemanse, N. J., Both, C., Drent, P. J., & Tinbergen, J. M. (2004). Fitness consequences of avian personalities in a fluctuating environment. Proceedings. Biological Sciences / The Royal Society, 271(1541), 847–852. 10.1098/rspb.2004.2680

Dingemanse, N. J., Both, C., Drent, P., Van Oers, K., & Van Noordwijk, A. J. (2002). Repeatability and heritability of exploratory behaviour in great tits from the wild. Animal Behaviour, 64(6), 929–938. 10.1006/anbe.2002.2006

Dochtermann, N. A. (2023). The role of plasticity, trade-offs, and feedbacks in shaping behavioral correlations. Behavioral Ecology, 34(6), 913–918. 10.1093/beheco/arad056

Durier, V., & Rivault, C. (2001). Effects of spatial knowledge and feeding experience on foraging choices in German cockroaches. Animal Behaviour, 62(4), 681–688. 10.1006/anbe.2001.1807

Farine, D. R. (2013). Animal social network inference and permutations for ecologists in R using asnipe. Methods in Ecology and Evolution, 4(12), 1187–1194. 10.1111/2041-210X.12121

Farine, D. R., & Whitehead, H. (2015). Constructing, conducting, and interpreting animal social network analysis. The Journal of Animal Ecology, 84(5), 1144–1163. 10.1111/1365-2656.12418

Farrar, B. G., Voudouris, K., & Clayton, N. S. (2021). Replications, Comparisons, Sampling and the Problem of Representativeness in Animal Cognition Research. Animal Behavior and Cognition, 8(2), 273–295. 10.26451/abc.08.02.14.2021

Fisher, D. N. (2023). Direct and indirect phenotypic effects on sociability indicate potential to evolve. Journal of Evolutionary Biology, 36(1), 209–220. 10.1111/jeb.14110

Freeman, L. C. (1978). Centrality in social networks conceptual clarification. Social Networks, 1(3), 215–239. 10.1016/0378-8733(78)90021-7

Gaona-Gordillo, I., Holtmann, B., Mouchet, A., Hutfluss, A., Sánchez-Tójar, A., & Dingemanse, N. J. (2023). Are animal personality, body condition, physiology and structural size integrated? A comparison of species, populations and sexes, and the value of study replication. Journal of Animal Ecology, 92(9), 1707–1718. 10.1111/1365-2656.13966

Godfrey, S. S., Sih, A., & Bull, C. M. (2013). The response of a sleepy lizard social network to altered ecological conditions. Animal Behaviour, 86(4), 763–772. 10.1016/j.anbehav.2013.07.016

Hadfield, J. D. (2010). MCMC methods for multi-response generalized linear mixed models: The MCMCglmm R package. Journal of Statistical Software, 33(2), 1–22.

Holand, Ø., Gjøstein, H., Losvar, A., Kumpula, J., Smith, M. E., Røed, K. H., Nieminen, M., & Weladji, R. B. (2004). Social rank in female reindeer (Rangifer tarandus): Effects of body mass, antler size and age. Journal of Zoology, 263(4), 365–372. 10.1017/S0952836904005382

Kesel, A. B., Martin, A., & Hoffmann, F. (2009). Quantifying the Landing Reaction of Cockroaches (Final Report 08/6302; 21958/08/NL/CBI, p. 56). European Space Agency.

Korsten, P., van Overveld, T., Adriaensen, F., & Matthysen, E. (2013). Genetic integration of local dispersal and exploratory behaviour in a wild bird. Nature Communications, 4(1), 2362. 10.1038/ncomms3362

Kralj-Fišer, S., & Schuett, W. (2014). Studying personality variation in invertebrates: Why bother? Animal Behaviour, 91, 41–52. 10.1016/j.anbehav.2014.02.016

Krause, J., Croft, D. P., & James, R. (2007). Social network theory in the behavioural sciences: Potential applications. Behavioral Ecology and Sociobiology, 62(1), 15–27. 10.1007/s00265-007-0445-8

Krause, J., James, R., Franks, D. W., & Croft, D. P. (2014). Animal Social Networks. Oxford University Press. http://ukcatalogue.oup.com/product/9780199679058.do

Lihoreau, M., & Rivault, C. (2010). German cockroach males maximize their inclusive fitness by avoiding mating with kin. Animal Behaviour, 80(2), 303–309. 10.1016/j.anbehav.2010.05.011

Lipton, J., Kleemann, G., Ghosh, R., Lints, R., & Emmons, S. W. (2004). Mate searching in Caenorhabditis elegans: A genetic model for sex drive in a simple invertebrate. Journal of Neuroscience, 24(34), 7427–7434. Scopus. 10.1523/JNEUROSCI.1746-04.2004

Louca, V., Lindsay, S. W., & Lucas, M. C. (2009). Factors triggering floodplain fish emigration: Importance of fish density and food availability. Ecology of Freshwater Fish, 18(1), 60–64. 10.1111/j.1600-0633.2008.00323.x

Luttbeg, B., & Sih, A. (2010). Risk, resources and state-dependent adaptive behavioural syndromes. Philosophical Transactions of the Royal Society of London. Series B, Biological Sciences, 365(1560), 3977–3990. 10.1098/rstb.2010.0207

Marjamäki, P. H., Contasti, A. L., Coulson, T. N., & McLoughlin, P. D. (2013). Local density and group size interacts with age and sex to determine direction and rate of social dispersal in a polygynous mammal. Ecology and Evolution, 3(9), 3073–3082. 10.1002/ece3.694

Opsahl, T. (2009). Structure and Evolution of Weighted Networks. University of London (Queen Mary College).

Opsahl, T., Agneessens, F., & Skvoretz, J. (2010). Node centrality in weighted networks: Generalizing degree and shortest paths. Social Networks, 32(3), 245–251. 10.1016/j.socnet.2010.03.006

Opsahl, T., & Panzarasa, P. (2009). Clustering in weighted networks. Social Networks, 31(2), 155–163. 10.1016/j.socnet.2009.02.002

Planas-Sitja, I., & Deneubourg, J. L. (2018). The role of personality variation, plasticity and social facilitation in cockroach aggregation. Biology Open, 7(12). 10.1242/BIO.036582/2072

Réale, D., Reader, S. M., Sol, D., McDougall, P. T., & Dingemanse, N. J. (2007). Integrating animal temperament within ecology and evolution. Biological Reviews, 82(2), 291–318. 10.1111/j.1469-185X.2007.00010.x

Ross, M. H., & Tignor, K. R. (1985). Response of german cockroaches to a dispersant emitted by adult females. Entomologia Experimentalis et Applicata, 39(1), 15–20. 10.1111/j.1570-7458.1985.tb03537.x

Sabol, A. C., Lambert, C. T., Keane, B., Solomon, N. G., & Dantzer, B. (2020). How does individual variation in sociality influence fitness in prairie voles? Animal Behaviour, 163, 39–49. 10.1016/j.anbehav.2020.02.009

Schielzeth, H. (2010). Simple means to improve the interpretability of regression coefficients. Methods in Ecology and Evolution, 1(2), 103–113. 10.1111/j.2041-210X.2010.00012.x

Schuett, W., Laaksonen, J., & Laaksonen, T. (2012). Prospecting at conspecific nests and exploration in a novel environment are associated with reproductive success in the jackdaw. Behavioral Ecology and Sociobiology, 66(9), 1341–1350. 10.1007/s00265-012-1389-1

Smith, B. R., & Blumstein, D. T. (2008). Fitness consequences of personality: A meta-analysis. Behavioral Ecology, 19(2), 448–455. 10.1093/beheco/arm144

Snijders, L., van Rooij, E. P., Burt, J. M., Hinde, C. A., van Oers, K., & Naguib, M. (2014). Social networking in territorial great tits: Slow explorers have the least central social network positions. Animal Behaviour, 98, 95–102. 10.1016/j.anbehav.2014.09.029

Sosa, S., Sueur, C., & Puga-Gonzalez, I. (2021). Network measures in animal social network analysis: Their strengths, limits, interpretations and uses. Methods in Ecology and Evolution, 12(1), 10–21. 10.1111/2041-210X.13366

Stahl, J., Tolsma, P. H., Loonen, M. J. J. E., & Drent, R. H. (2001). Subordinates explore but dominants profit: Resource competition in high Arctic barnacle goose flocks. Animal Behaviour, 61(1), 257–264. 10.1006/anbe.2000.1564

Stanley, C. R., Liddiard Williams, H., & Preziosi, R. F. (2018). Female clustering in cockroach aggregations-A case of social niche construction? Ethology, 124(10), 706–718. 10.1111/eth.12799

Stanley, C. R., Mettke-Hofmann, C., & Preziosi, R. F. (2017). Personality in the cockroach Diploptera punctata: Evidence for stability across developmental stages despite age effects on boldness. PLoS ONE, 12(5). 10.1371/journal.pone.0176564

Sueur, C., Petit, O., De Marco, A., Jacobs, A. T., Watanabe, K., & Thierry, B. (2011). A comparative network analysis of social style in macaques. Animal Behaviour, 82(4), 845–852. 10.1016/j.anbehav.2011.07.020

Székely, T. (Tamás), Moore, A. J. (Allen J., & Komdeur, J. (2010). Social behaviour: Genes, ecology and evolution. Cambridge University Press. https://www.cambridge.org/gb/academic/subjects/life-sciences/animal-behaviour/social-behaviour-genes-ecology-and-evolution?format=PB&isbn=9780521709620

Tignor, K. R., & Ross, M. H. (1987). Effect of starvation on secretion of a dispersal pheromone by female german cockroaches. Entomologia Experimentalis et Applicata, 45(3), 245–249. 10.1111/j.1570-7458.1987.tb01090.x

Turner, J. W., Bills, P. S., & Holekamp, K. E. (2018). Ontogenetic change in determinants of social network position in the spotted hyena. Behavioral Ecology and Sociobiology, 72(1). 10.1007/s00265-017-2426-x

van Overveld, T., Careau, V., Adriaensen, F., & Matthysen, E. (2014). Seasonal- and sex-specific correlations between dispersal and exploratory behaviour in the great tit. Oecologia, 174(1), 109–120. 10.1007/s00442-013-2762-0

Verbeek, M. E. M., Boon, A., & Drent, P. J. (1996). Exploration, Aggressive Behaviour and Dominance in Pair-Wise Confrontations of Juvenile Male Great Tits. Behaviour, 133(11–12), 945–963. 10.1163/156853996X00314

Verbeek, M. E. M., Drent, P., & Wiepkema, P. (1994). Consistent individual differences in early exploratory behaviour of male great tits. Animal Behaviour, 48(5), 1113–1121.

Wey, T. W., Blumstein, D. T., Shen, W., & Jordán, F. (2008). Social network analysis of animal behaviour: A promising tool for the study of sociality. Animal Behaviour, 75(2), 333–344. 10.1016/j.anbehav.2007.06.020

Whitehead, H., & Dufault, S. (1999). Techniques for Analyzing Vertebrate Social Structure Using Identified Individuals: Review and Recommendations. Advances in the Study of Behavior, 28. http://www.sciencedirect.com/science/article/pii/S0065345408602156

Yip, E. C., Smith, D. R., & Lubin, Y. (2021). Causes of plasticity and consistency of dispersal behaviour in a group-living spider. Animal Behaviour, 175, 99–109. 10.1016/j.anbehav.2021.02.019

