## Supplementary materials for "Cockroach social network position does not influence exploration tendency"

Comparing networks built from 6 or 10 observations.

In order to compare network measures from social networks built from either six or ten observations, we used only data from block 2. In this block, ten observations of social networks were made, but only the first six were used in the analysis we present in the main text, to allow better comparison with block 1. Here we built social networks for each group in bock 2 with all ten observations and quantified strength, clustering coefficient, and closeness, to compare with the same measures calculated based on networks using only the first size observations.

Strength was very strongly positively correlated between the two networks (Pearson correlation, r = 0.911, t = 21.856, df = 98, p < 0.001; Fig. S1a). Clustering coefficient was moderately positively correlated between the two networks (Pearson correlation, r = 0.569, t = 6.848, df = 98, p < 0.001; Fig. S1b). Closeness was strongly positively correlated between the two networks (Pearson correlation, r = 0.761, t = 11.623, df = 98, p < 0.001; Fig. S1c).

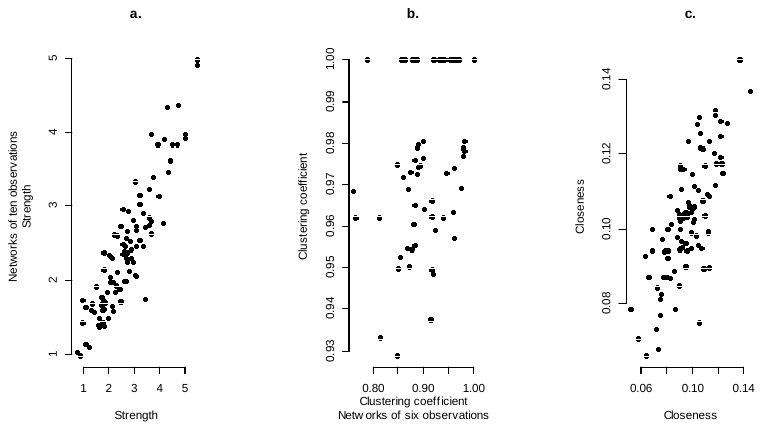

Figure S1. Plots of network measures calculated on six observations (x axes) and ten observations (y axes): a) Strength, b) Clustering coefficient, c) Closeness.

Results of analysis with those that did not explore in the group setting set the maximum time of 1440 seconds

To test for how sensitive our analysis was to the decision to not give an exploration score to those individuals who did not explore, we repeated the analysis with those individuals instead receiving the maximum exploration time of 1440 seconds (the model is otherwise identical). The results are very similar to the analysis presented in the main text, the only differences are that a positive relationship between mass and exploration time, suggesting a negative effect of mass on exploration tendency, was now apparent in males (posterior distribution mode = 1.242, posterior distribution mean = 1.254, 95% credible intervals = 0.012 to 2.5161, pMCMC = 0.050). Full results are given in Table S2.

Supplementary tables

Table S1. Residual variances (“*trait name* Residual Variance” and covariances (*trait name 1 – trait name 2*”) for the analysis reported in the main text. We give posterior distribution mode, mean, and its lower and upper 95% credible intervals (CrI).

| Variable | Posterior mode | Posterior mean | Lower 95% CrI | Upper 95% CrI |
| --- | --- | --- | --- | --- |
| Exploration alone Residual Variance | 0.8944 | 0.9102 | 0.6683 | 1.1750 |
| Exploration group Residual Variance | 1.6750 | 1.7892 | 1.4332 | 2.2041 |
| Strength Residual Variance | 0.3935 | 0.4024 | 0.3239 | 0.4864 |
| Closeness Residual Variance | 0.0003 | 0.0003 | 0.0002 | 0.0004 |
| Clustering Coefficient Residual Variance | 0.0011 | 0.0011 | 0.0009 | 0.0013 |
| Exploration alone - Exploration group | -0.1264 | -0.1067 | -0.3253 | 0.1031 |
| Exploration alone - Strength | 0.0462 | 0.0338 | -0.0663 | 0.1303 |
| Exploration alone - Closeness | 0.0013 | 0.0011 | -0.0016 | 0.0037 |
| Exploration alone - Clustering Coefficient | -0.0006 | -0.0004 | -0.0060 | 0.0048 |
| Exploration group - Strength | -0.0365 | -0.0638 | -0.1995 | 0.0597 |
| Exploration group - Closeness | -0.0020 | -0.0019 | -0.0053 | 0.0017 |
| Exploration group - Clustering Coefficient | -0.0040 | -0.0043 | -0.0111 | 0.0024 |
| **Strength - Closeness** | **0.0100** | **0.0104** | **0.0082** | **0.0125** |
| Strength - Clustering Coefficient | 0.0000 | 0.0001 | -0.0028 | 0.0032 |
| Closeness - Clustering Coefficient | 0.0000 | 0.0000 | -0.0001 | 0.0001 |

Table S2. Ful model results for the analysis where non exploring individuals were given the maximum score of 1440 seconds. For each of the five traits (exploration alone, exploration group, strength, closeness, and clustering coefficient) we give the fixed effects (intercept, the effect of mass, the effect of sex [females are the default, so this effect represents how males deviate from females], the mass:sex interaction [again, as females are the default this effect indicates how the effect of mass on male traits differs from that for females, which is indicated by the main effect of mass], the effect of block [block 1 is the default, and so this effect indicates how block 2 differs]), the random effect of group, and the residual variances and covariances (indicated where two trait names are given in the “Variable” column). As group is a random effect, and for the residual (co)variances) we only report the posterior mode, mean, and lower and upper 95% credible intervals (CrI), while as the remaining terms are fixed effects we also report the pMCMC value, which represents two times the probability the effect is either greater or smaller than zero (whichever is smaller), and can be interpreted in a similar way to traditional (frequentist) p-values. Terms where the 95% CrIs do not overlap with zero are highlighted in bold.

| Variable | Posterior mode | Posterior mean | Lower 95% CrI | Upper 95% CrI | pMCMC | Effect |
| --- | --- | --- | --- | --- | --- | --- |
| Intercept - Exploration alone | 1.0941 | 1.0843 | 0.6437 | 1.5015 | 0.0002 | fixed |
| Intercept - Exploration group | 5.0127 | 4.8968 | 4.1037 | 5.6795 | 0.0002 | fixed |
| Intercept - Strength | 2.3158 | 2.3445 | 1.8551 | 2.8650 | 0.0002 | fixed |
| Intercept - Closeness | 0.0945 | 0.0942 | 0.0828 | 0.1062 | 0.0002 | fixed |
| Intercept - Clustering Coefficient | 0.9441 | 0.9330 | 0.8857 | 0.9793 | 0.0002 | fixed |
| Mass – Exploration alone | 0.0909 | 0.0970 | -0.3147 | 0.4754 | 0.6260 | fixed |
| Mass – Exploration group | 0.2731 | 0.2318 | -0.2699 | 0.7189 | 0.3572 | fixed |
| Mass - Strength | 0.0792 | 0.0706 | -0.1566 | 0.3012 | 0.5404 | fixed |
| Mass - Closeness | 0.0032 | 0.0024 | -0.0038 | 0.0087 | 0.4572 | fixed |
| Mass - Clustering Coefficient | 0.0020 | -0.0007 | -0.0120 | 0.0114 | 0.9092 | fixed |
| Sex – Exploration alone | -0.0985 | -0.1218 | -1.1044 | 0.7447 | 0.7948 | fixed |
| Sex – Exploration group | 0.5365 | 0.5980 | -0.7172 | 1.9739 | 0.4120 | fixed |
| Sex - Strength | -0.7144 | -0.6101 | -1.3868 | 0.1181 | 0.1112 | fixed |
| Sex - Closeness | -0.0055 | -0.0012 | -0.0197 | 0.0172 | 0.8996 | fixed |
| Sex - Clustering Coefficient | -0.0480 | -0.0450 | -0.1037 | 0.0165 | 0.1336 | fixed |
| **Block – Exploration alone** | 1.0470 | 1.0042 | 0.6628 | 1.3092 | 0.0002 | fixed |
| Block – Exploration group | 0.2815 | 0.3676 | -0.3954 | 1.1948 | 0.3532 | fixed |
| **Block - Strength** | 0.6604 | 0.6486 | 0.0771 | 1.2103 | 0.0244 | fixed |
| Block - Closeness | -0.0007 | 0.0004 | -0.0116 | 0.0133 | 0.9528 | fixed |
| Block - Clustering Coefficient | 0.0194 | 0.0219 | -0.0327 | 0.0770 | 0.3944 | fixed |
| Mass:Sex – Exploration alone | -0.3287 | -0.2090 | -1.1657 | 0.7156 | 0.6652 | fixed |
| **Mass:Sex – Exploration group** | 1.2415 | 1.2535 | 0.0115 | 2.5162 | 0.0504 | fixed |
| Mass:Sex - Strength | -0.1258 | -0.1030 | -0.6724 | 0.4725 | 0.7256 | fixed |
| Mass:Sex - Closeness | -0.0041 | -0.0050 | -0.0204 | 0.0109 | 0.5404 | fixed |
| Mass:Sex - Clustering Coefficient | -0.0135 | -0.0093 | -0.0399 | 0.0209 | 0.5524 | fixed |
| Group – Exploration alone | 0.0013 | 0.0123 | 0.0002 | 0.0477 | NA | random |
| Group – Exploration group | 0.5107 | 0.6571 | 0.1493 | 1.2813 | NA | random |
| Group - Strength | 0.2863 | 0.3884 | 0.1498 | 0.6807 | NA | random |
| Group - Closeness | 0.0001 | 0.0002 | 0.0001 | 0.0003 | NA | random |
| Group - Clustering Coefficient | 0.0034 | 0.0038 | 0.0016 | 0.0066 | NA | random |
| Exploration alone Residual | 0.8908 | 0.9116 | 0.6639 | 1.1657 | NA | residual |
| Exploration alone – Exploration group | -0.1731 | -0.1555 | -0.3728 | 0.0550 | NA | residual |
| Exploration alone - Strength | 0.0393 | 0.0342 | -0.0665 | 0.1314 | NA | residual |
| Exploration alone - Closeness | 0.0009 | 0.0011 | -0.0016 | 0.0038 | NA | residual |
| Exploration alone – Clustering coefficient | -0.0009 | -0.0004 | -0.0056 | 0.0051 | NA | residual |
| Exploration group Residual | 1.8818 | 1.9319 | 1.5604 | 2.3383 | NA | residual |
| Exploration group - Strength | -0.0824 | -0.0815 | -0.2126 | 0.0444 | NA | residual |
| Exploration group - Closeness | -0.0023 | -0.0025 | -0.0059 | 0.0011 | NA | residual |
| Exploration group - Clustering | -0.0025 | -0.0037 | -0.0105 | 0.0031 | NA | residual |
| Strength Residual | 0.3855 | 0.4021 | 0.3224 | 0.4846 | NA | residual |
| **Strength - Closeness** | **0.0103** | **0.0104** | **0.0083** | **0.0127** | NA | residual |
| Strength – Clustering coefficient | 0.0002 | 0.0001 | -0.0029 | 0.0031 | NA | residual |
| Closeness Residual | 0.0003 | 0.0003 | 0.0002 | 0.0004 | NA | residual |
| Closeness – Clustering coefficient | 0.0000 | 0.0000 | -0.0001 | 0.0001 | NA | residual |
| Clustering coefficient Residual | 0.0011 | 0.0011 | 0.0009 | 0.0013 | NA | residual |
